# Clinical and *In Situ* Characterization of Hepatitis Delta Virus in Sjogren’s Syndrome

**DOI:** 10.1101/2022.12.03.518993

**Authors:** Matthew C. Hesterman, Summer V. Furrer, Braden S. Fallon, Melodie L. Weller

## Abstract

Hepatitis delta virus (HDV) has been detected in the minor salivary gland (MSG) tissue of Sjogren’s Syndrome (SjS) patients in the absence of an HBV co-infection. HDV antigen expression was previously shown to trigger an SjS-like phenotype in vivo, demonstrating a cause-and-effect association. We hypothesize that if HDV regulates SjS development, then HDV profiles may correlate with disease manifestations. This retrospective study characterized HDV in a cohort of 48 SjS MSG between 2014-2021. Analyses of HDV antigen (HDAg) expression, including cell type and subcellular localization, *in situ* hybridization of HDV RNA, and comparative analyses with associated SjS and viral hepatitis clinical features were conducted. HDAg was detected in MSG acinar, ductal, and adipose cells. HDAg localized with nuclei and mitochondria. HDV genomic RNA localized to the nucleus. A significant negative correlation was noted between HDAg intensity and focal lymphocytic inflammation. No significant associations were detected between MSG-localized HDAg and liver enzymes, or an evident HBV co-infection. This study has identified a non-hepatic reservoir for chronic HDV persistence in SjS-affected MSG and a unique mitochondrial localization for HDV antigen. Detection of non-hepatic HDV-mediated disease in the absence of an evident current or past HBV co-infection warrants further investigation.

## Introduction

Sjogren’s syndrome (SjS) or Sjogren’s Disease (SjD) is a non-communicable, chronic autoimmune disease afflicting ≤0.1% of the population in the United States (1–3). SjS is currently diagnosed by the detection of decreased tear and/or saliva production, development of autoantibodies (anti-SSA/Ro, Anti-SSB/La), and focal lymphocytic inflammation within the salivary gland tissue and in connection with significant extraglandular manifestations (4–6). The etiology of this female predominant disease is attributed to a combination of genetic susceptibility factors, hormone profiles and/or chronic pathogen exposure (7–10). Previously, hepatitis delta virus (HDV) was detected in SjS salivary gland tissue and shown to trigger an SjS-like phenotype *in vivo* (11). This current retrospective study builds upon the discovery of HDV in SjS patients and aims to characterize the clinical and *in situ* features of HDV within an Utahn SjS cohort.

Hepatitis delta virus is a rare infectious disease estimated to impact 12-72 million people worldwide (12– 14). As a member of the Kolmioviridae family, HDV is the only known viral satellite RNA (sat-RNA) to infect humans (Figure 1). This ∼1700nt circular ssRNA sat-RNA produces two antigens: the small HDAg (S-HDAg) and large HDAg (L-HDAg) from a single open reading frame (15–17). A helper virus is required for packaging and transmission (ex. hepatitis B virus (HBV)) and the tissue tropism of HDV is dictated by the tropism of the helper virus (18). Once intracellular, HDV utilizes the S-HDAg and host cellular proteins for replication and can persist in the absence of an active helper virus co-infection (19). The L-HDAg facilitates packaging of the ribonucleoprotein complex into the helper virus particle (HBV) (20). Hepatic HDV co-infection with HBV is associated with enhanced symptoms of viral hepatitis, increased risk of liver decompensation and hepatocellular carcinoma (21,22). Since the discovery of HDV in the 1970s (23), this unique pathogen has been extensively studied in the context of viral hepatitis. The recent discovery of HDV infection of non-hepatic tissue and associated development of exocrine gland dysfunction suggests that there may be more to this sat-RNA than previously defined (11).

**Figure 1.**
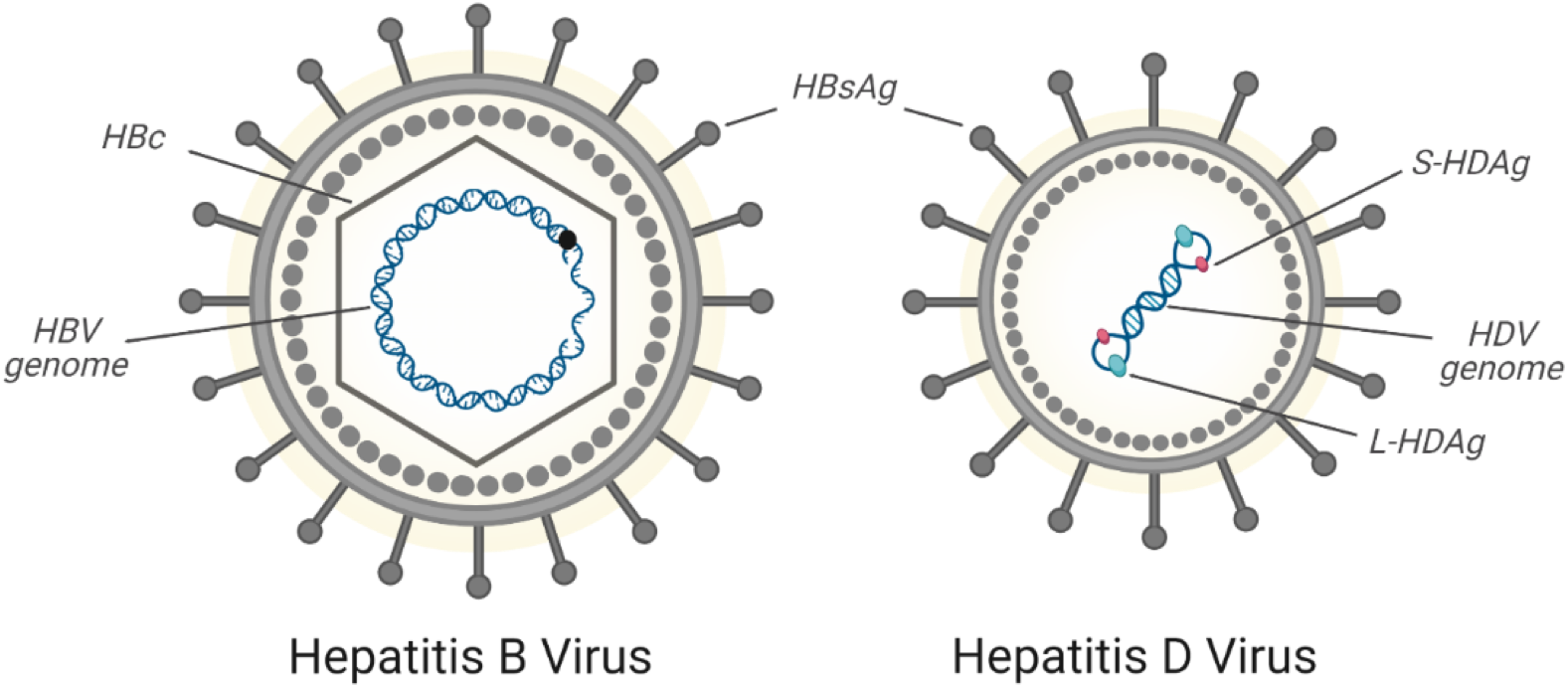
Hepatitis delta virus (HDV) and Hepatitis B Virus (HBV) Overview. HDV is the only satellite RNA known to infect humans and requires a helper virus (ex. hepatitis B virus (HBV)) for packaging and transmission. The cellular tropism of HDV-containing particles is dictated by the helper virus’ encapsulation proteins. Created with BioRender.com.

HDV sequence and antigen were previously detected in minor salivary gland (MSG) tissue of Sjogren’s syndrome patients (11). A current or prior HBV co-infection was not detected in patients with detectible HDV within salivary gland tissue. Additionally, expression of HDV antigens in a murine model triggered the development of a complete SjS-like phenotype *in vivo*. S-HDAg expression triggered the development of autoantibodies (anti-SSA/Ro, anti-SSB/La) and decreased stimulated saliva flow in the murine model. Mice expressing both the S-HDAg and L-HDAg developed significant focal inflammation within the salivary gland tissue and anti-SSA/Ro antibodies. These findings support a cause-and-effect relationship between HDV antigen expression and the SjS phenotype.

The presence and impact of HDV localized outside of hepatic tissue has not been well defined. Retrospective analysis of an SjS patient cohort was performed to evaluate HDV antigen and sequence localization in labial MSG biopsies. The HDV profiles within the Utahn SjS patient cohort were then used in comparative analyses with Sjogren’s syndrome diagnostic criteria, autoimmune disease co-morbidities, liver enzymes, and viral hepatitis tests. Through this study, HDV was found to localize within multiple salivary gland cell types and with a unique subcellular localization pattern. Together, this study presents essential information on HDV co-localization within SjS minor salivary gland tissue and identifies SjS clinical parameters significantly correlating with HDV antigen expression levels.

## Results

### Sjogren’s syndrome patient demographics and clinical characterization

This study characterized hepatitis delta virus (HDV) and associated clinical parameters in the Utahn Sjogren’s syndrome (SjS) patient cohort through retrospective analyses. Forty-eight (48) archived MSG biopsies from SjS patients were analyzed (Table 1). In line with the established female predominance of Sjogren’s syndrome, the study cohort consisted of 91.7% females and 8.3% males with an average age of 49.9 ± 13.2 years at Sjogren’s syndrome diagnosis. Lymphocytic focus scores ≥1 per 4mm^2^ area were observed in 33/48 (68.8%) of minor salivary gland biopsies analyzed. Antibody testing identified positivity rates of 15/31 (48.4%) for antinuclear antibodies (ANA), 12/44 (27.3%) for anti-SSA/Ro, 1/43 (2.1%) for anti-SSB/La, and 5/33 (15.2%) for rheumatoid factor (RF). No evidence of hypocomplementemia of C3 and C4 were noted in the SjS cohort as has been reported in prior SjS studies (Supplemental Table 1) (24). Abnormal (high) C3 and C4 levels were identified in 2/21 (9.1%) and 4/22 (18.2%) patients, respectively (Supplemental Table 1).

**Table 1.** Sjogren’s Syndrome patient demographics and characterization of clinical features.

|  | **Sjogren's**<br>**Syndrome**<br>**(n = 48)** |
| --- | --- |
| **Sex** |  |
| Female | 44 (91.7%) |
| Male | 4 (8.3%) |
| **Race** |  |
| Asian | 0 (0.0%) |
| American Indian or Alaskan Native | 0 (0.0%) |
| Black or African American | 0 (0.0%) |
| Native Hawaiian or Pacific Islander | 0 (0.0%) |
| White | 45 (93.8%) |
| Other | 3 (6.3%) |
| **Ethnicity** |  |
| Hispanic/ Latino | 5 (10.4%) |
| Not Hispanic/ Latino | 43 (89.6%) |
| **Age (M ± SD)** | 49.9 ± 13.2 |
| **Age Group** |  |
| 0 – 19 | 0 (0.0%) |
| 20 – 39 | 10 (20.8%) |
| 40 – 59 | 25 (52.1%) |
| 60+ | 13 (27.1%) |
| **Foci/4mm<sup>2</sup>** | 1.5 ± 1.9 |
| ≥1 | 33 (68.8%) |
| <1 | 15 (31.3%) |
| **Autoantibodies** |  |
| ANA (+) | 15 / 31 (48.4%) |
| Anti-SSA (+) | 12 / 44 (27.3%) |
| Anti-SSB (+) | 1 / 43 (2.1%) |
| RF (+) | 5 / 33 (15.2%) |

Abbreviations: M = mean, SD = standard deviation, RF = rheumatoid factor, ANA = antinuclear antibody

## HDV antigens detected in SjS minor salivary gland tissue

### HDAg detected in Acinar and Ductal cells

Labial MSG biopsies were evaluated for the cellular localization of HDAg. Cellular localization of HDAg was performed using immunohistochemical staining of aquaporin 5 (AQP5) and cytokeratin 7 (CK7) to detect acinar and ductal cells, respectively. As detailed in Figure 2A, HDAg localized within both acinar and ductal cells of the labial MSG tissues. Additionally, HDAg co-localized with the nuclear region of adipocyte cells within the connective tissue (Supplemental Figure 1). HDAg expression was not evident in lymphocytic foci (data not shown).

**Figure 2.**
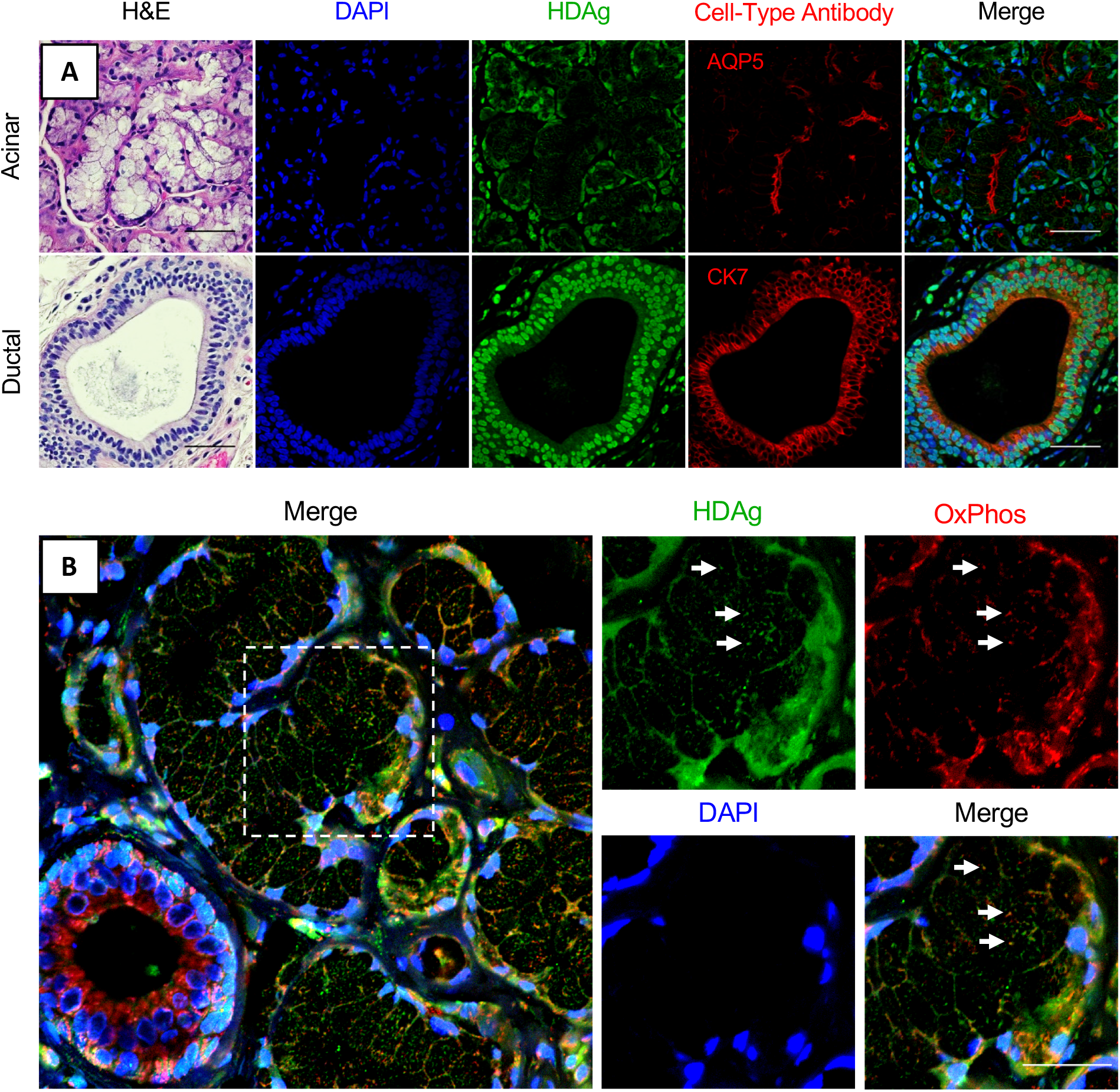
Cellular and subcellular localization of HDAg in SjS labial minor salivary glands. **A**) Cellular localization patterns of HDAg were identified through immunohistochemical staining of formalin-fixed paraffin-embedded (FFPE) minor labial salivary gland sections. Cell nuclei were stained with DAPI (blue); anti-HDAg antibody was used for immunostaining of HDAg (green). Antibodies for AQP5 and CK7 were used for identification of acinar and ductal cells, respectively (red). Scale bars = 50μm. **B**) Subcellular co-localization of HDAg with mitochondria was observed. Cell nuclei were stained with DAPI (blue); anti-HDAg antibody was used for immunostaining of HDAg (green). OxPhos Human Antibody Cocktail was used for identification of mitochondria (red). Scale bars = 25μm.

### HDAg co-localize with the nucleus and mitochondria

Labial minor salivary gland tissues were evaluated for subcellular localization patterns of HDAg. Immunohistochemical analysis of labial MSG FFPE tissues with DAPI and OxPhos antibody cocktail was performed to further characterize subcellular localization of HDAg in the nucleus and mitochondria, respectively. As detailed in Figure 2A-B, HDAg was detected in the nucleus of both acinar and ductal cells. HDAg in adipocytes presented with a similar nuclear localization pattern (Supplemental Figure 1). HDAg was also shown to co-localize with the mitochondria in both acinar and ductal cell types (Figure 2B).

### Nuclear localized HDV sequence detected in SjS minor salivary gland tissue

*In situ* hybridization was performed to detect HDV genomic and anti-genomic RNA sequence utilizing custom RNAScope® probes. As detailed in Figure 3, HDV genomic RNA was detected and localized within the cell nuclei (Figure 3A). HDV RNA did not co-localize with mitochondrial RNA as measured by an RNAscope® probe detecting COX I RNA (Figure 3B). Probes against the anti-genomic HDV sequence did not detect sequence within the labial MSG tissues analyzed (data not shown). This lack of antigenomic sequence may be due to the lower copy number of antigenomic sequences compared to genomic sequences (1:15) that are present during the HDV replication cycle (25). Nuclear localization of HDV sequence has previously been associated with active replication of the viral genome (26).

**Figure 3.**
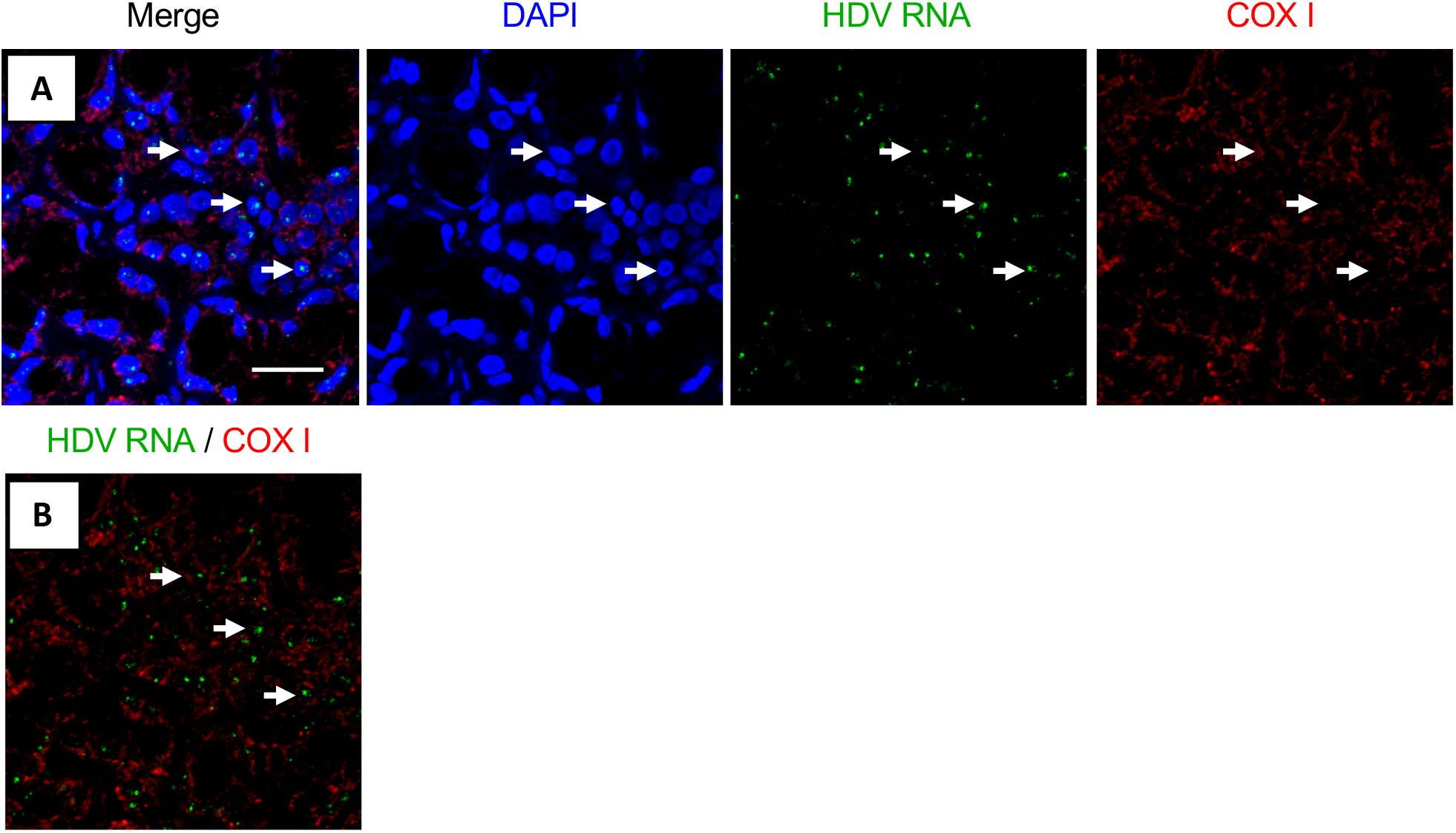
Subcellular colocalization of HDV genomic RNA with nucleus in salivary gland biopsies of SjS patients. **A**) Merged overview of MSG tissue with cell nuclei (blue), HDV RNA (green), and COX I RNA (red). **B**) Merged overview of in situ hybridization of probes for HDV RNA and COX I demonstrating lack of colocalization between HDV RNA and mitochondrial RNA. Arrows indicate examples of colocalization of HDV RNA with cell nuclei; scale bar = 25μm

## HDV-SjS Clinical Parameters

### HDAg intensity negatively correlates with lymphocytic foci

Comparative analyses were performed between patients’ HDV antigen intensity profile within MSG biopsies and clinical parameters of SjS diagnosis, including focal lymphocytic inflammation, autoantibody profile, and demographics. HDAg intensity was classified into 3 categories (high, moderate, low intensity) as outlined in the methods section and Supplemental Figure 2. There were no significant differences in HDV intensity based on race or ethnicity (Table 2). A significant difference was noted between the median age of patients in each HDAg intensity group, with the median age of the high HDAg intensity group being 49.5 *±* 10.4, moderate intensity group being 42.1 ± 13.3, and low intensity group being 57.2 ± 12.1 (ANOVA, p=0.004) (Table 2). Additionally, a significant difference between HDAg intensity groupings and age groups was observed with increased HDAg intensity in younger age groups (age group 20-39) and lower HDAg intensity in 60+ age group (p=0.03) (Table 2).

**Table 2.**
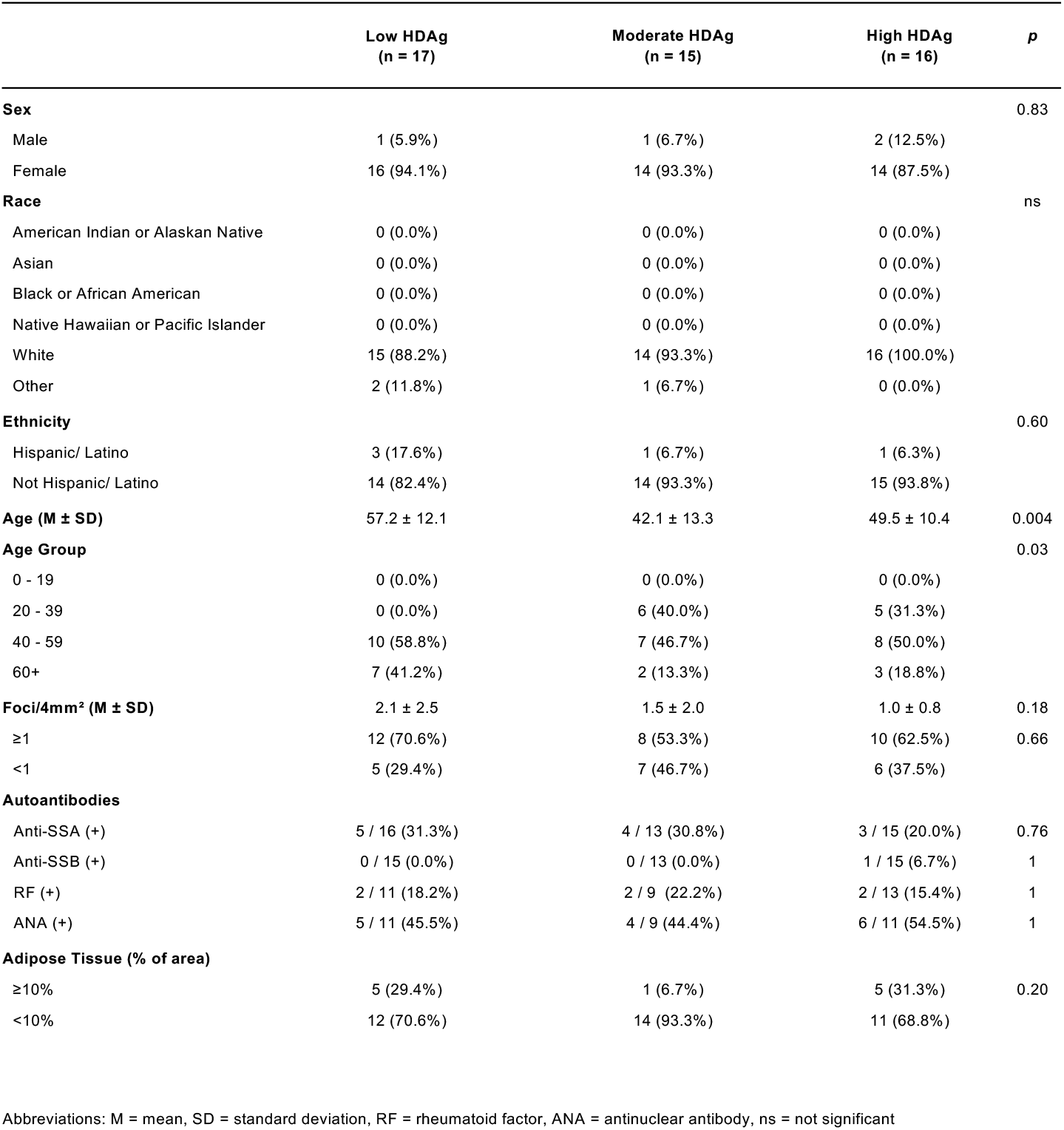
Patient demographics and primary Sjogren’s syndrome diagnostic data stratified by HDV antigen intensity level.

|  | Low HDAG<br>(n = 17) | Moderate HDAG<br>(n = 15) | High HDAG<br>(n = 16) | p |
| --- | --- | --- | --- | --- |
| **Sex** |  |  |  | 0.83 |
| Male | 1 (5.9%) | 1 (6.7%) | 2 (12.5%) |  |
| Female | 16 (94.1%) | 14 (93.3%) | 14 (87.5%) |  |
| **Race** |  |  |  | ns |
| American Indian or Alaskan Native | 0 (0.0%) | 0 (0.0%) | 0 (0.0%) |  |
| Asian | 0 (0.0%) | 0 (0.0%) | 0 (0.0%) |  |
| Black or African American | 0 (0.0%) | 0 (0.0%) | 0 (0.0%) |  |
| Native Hawaiian or Pacific Islander | 0 (0.0%) | 0 (0.0%) | 0 (0.0%) |  |
| White | 15 (88.2%) | 14 (93.3%) | 16 (100.0%) |  |
| Other | 2 (11.8%) | 1 (6.7%) | 0 (0.0%) |  |
| **Ethnicity** |  |  |  | 0.60 |
| Hispanic/ Latino | 3 (17.6%) | 1 (6.7%) | 1 (6.3%) |  |
| Not Hispanic/ Latino | 14 (82.4%) | 14 (93.3%) | 15 (93.8%) |  |
| **Age (M ± SD)** | 57.2 ± 12.1 | 42.1 ± 13.3 | 49.5 ± 10.4 | 0.004 |
| **Age Group** |  |  |  | 0.03 |
| 0 - 19 | 0 (0.0%) | 0 (0.0%) | 0 (0.0%) |  |
| 20 - 39 | 0 (0.0%) | 6 (40.0%) | 5 (31.3%) |  |
| 40 - 59 | 10 (58.8%) | 7 (46.7%) | 8 (50.0%) |  |
| 60+ | 7 (41.2%) | 2 (13.3%) | 3 (18.8%) |  |
| **Foci/4mm<sup>2</sup> (M ± SD)** | 2.1 ± 2.5 | 1.5 ± 2.0 | 1.0 ± 0.8 | 0.18 |
| ≥1 | 12 (70.6%) | 8 (53.3%) | 10 (62.5%) | 0.66 |
| <1 | 5 (29.4%) | 7 (46.7%) | 6 (37.5%) |  |
| **Autoantibodies** |  |  |  |  |
| Anti-SSA (+) | 5 / 16 (31.3%) | 4 / 13 (30.8%) | 3 / 15 (20.0%) | 0.76 |
| Anti-SSB (+) | 0 / 15 (0.0%) | 0 / 13 (0.0%) | 1 / 15 (6.7%) | 1 |
| RF (+) | 2 / 11 (18.2%) | 2 / 9 (22.2%) | 2 / 13 (15.4%) | 1 |
| ANA (+) | 5 / 11 (45.5%) | 4 / 9 (44.4%) | 6 / 11 (54.5%) | 1 |
| **Adipose Tissue (% of area)** |  |  |  |  |
| ≥10% | 5 (29.4%) | 1 (6.7%) | 5 (31.3%) | 0.20 |
| <10% | 12 (70.6%) | 14 (93.3%) | 11 (68.8%) |  |

Abbreviations: M = mean, SD = standard deviation, RF = rheumatoid factor, ANA = antinuclear antibody, ns = not significant

Evaluation of HDAg intensity within MSG identified multiple significant correlations with demographics and clinical features in the Utahn SjS cohort. Hepatitis delta virus antigen intensity negatively correlated with focus score (r=-0.30, p<0.05) (Figure 4A). No significant difference was noted in HDAg intensity for biopsies with focus scores of <1 or ≥1 foci/4mm^2^ (Figure 4B). A negative trend was observed between HDAg intensity and age, with HDAg intensity steadily declining with increasing age (Figure C-D). An increasing trend was observed between focus score and age groups (ns) (Figure 4E).

**Figure 4.**
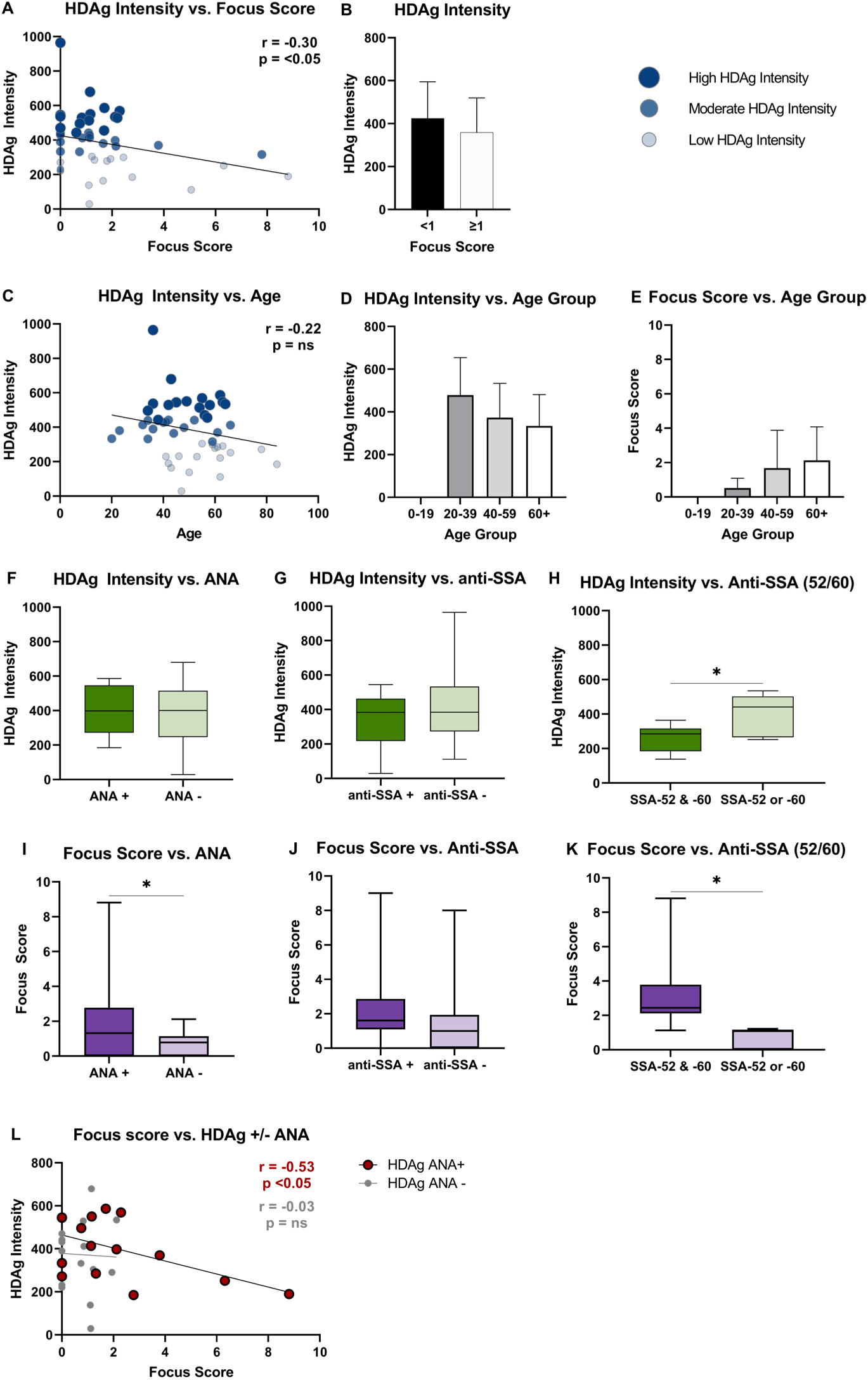
HDAg intensity comparisons and clinical characteristics correlation analysis. **A**) Pearson correlation analysis between HDAg fluorescence intensity of MSG biopsy samples and biopsy focus score, n = 48. **B**) Welch’s t-test comparison between HDAg fluorescence intensity and focus score grouped by focus score ≥1 and <1, n = 48. **C**) Pearson correlation between HDAg fluorescence intensity of MSG biospy samples and age n = 48. Age groups of biopsied patients were then compared with **D**) HDAg intensity and **E**) focus score, n = 48. Patients’ biopsies’ HDAg intensity was compared against **F**) anti-nuclear antibody testing data (n = 15) **G**) anti-SSA antibody (n = 12). **H**) Patients with SSA-52 and – 60 and patients with either SSA-52 or –60 were compared by HDAg intensity (n = 12). **I**) Patients’ biopsies’ focus scores were compared against **F**) anti-nuclear antibody testing data (n = 15) **G**) anti-SSA antibody (n = 12). K) Patients with SSA-52 and –60 and patients with either SSA-52 or –60 were compared by focus score (n = 12).**L**) Patients’ biopsies were grouped according to ANA test status and plotted according to HDAg intensity and focus score. Statistics were done in GraphPad Prism (**A, C** and **L**, Pearson correlation; **B, F**-**K**, Welch’s t-test; **D** and **E**, 1-way ANOVA with multiple comparison’s test). Bar graph data are shown as mean ± SD, scatterplots determine \**P* < 0.05.

Autoantibody profiles were evaluated relative to HDAg intensity. No significant differences were observed between HDAg intensity and the presence of anti-nuclear autoantibody (ANA) or anti-SSA (Figure 4F-G). Patients positive for either anti-SSA 52 or 60 exhibited elevated HDAg intensity levels compared to patients that were positive for both anti-SSA 52 and 60 (p < 0.05) (Figure 4H).

Autoantibody profiles were evaluated in relation to focal inflammation. Patients who tested positive for ANA had a significantly increased focus score compared to patients that were negative for ANA (p < 0.05) (Figure 4I). No difference in focus score was detected between patients who tested positive or negative for anti-SSA (Anti-SSA-52 or -60) (Figure 4J). In contrast to HDAg intensity, patients who were positive for both anti-SSA-52 and -60 had a significantly increased in focus score relative to patients that were positive for either anti-SSA-52 or -60 (p<0.05) (Figure 4K). In SjS patients that were ANA(+), HDAg intensity negatively correlated with focus score (Figure 4L). No correlation was observed between HDAg intensity and focus score in ANA(-) patients. Together, HDAg intensity negatively correlated with focus score and was decreased in patients that were positive for both anti-SSA-52 and anti-SSA-60.

Hepatitis delta virus antigen was detected in nuclei of adipose tissue (Supplemental Figure 1). Therefore, percent area of adipose tissue within MSG was evaluated for correlations with other clinical and demographic data. No significant differences were seen between the HDAg intensity and percentage of adipose tissue within the MSG tissue (Supplemental Figure 4). No significant correlations were observed between adipose area and focus score, age, sex, or auto-antibody profile (data not shown).

### Comorbidities in HDV(+) SjS patients

Evaluation of autoimmune and infectious disease comorbidities were performed with our defined Utahn Sjogren’s syndrome cohort (Table 3). Of the autoimmune diseases evaluated, SjS patients diagnosed with autoimmune thyroiditis and/or hypothyroidism were significantly more represented in the high HDAg intensity group relative to low and moderate HDAg intensity (p<0.006). Within the Utahn SjS cohort analyzed, 14.6% (7/48) of SjS patients and 37.5% (6/16) of SjS paitents with high HDAg intensity had a diagnosis of autoimmune thyroiditis or hypothyroidism (Table 3). Additionally, there was an increasing trend of myositis in the high HDAg intensity group (ns). No prior Hepatitis C Virus (HCV) or HBV diagnoses were detected in the SjS cohort exhibiting no significant difference in HDAg intensity. Other comorbidities (lupus, rheumatoid arthritis, multiple sclerosis, ankylosing spondylitis) exhibited no significant trends based on patients’ HDAg intensity group.

**Table 3.**
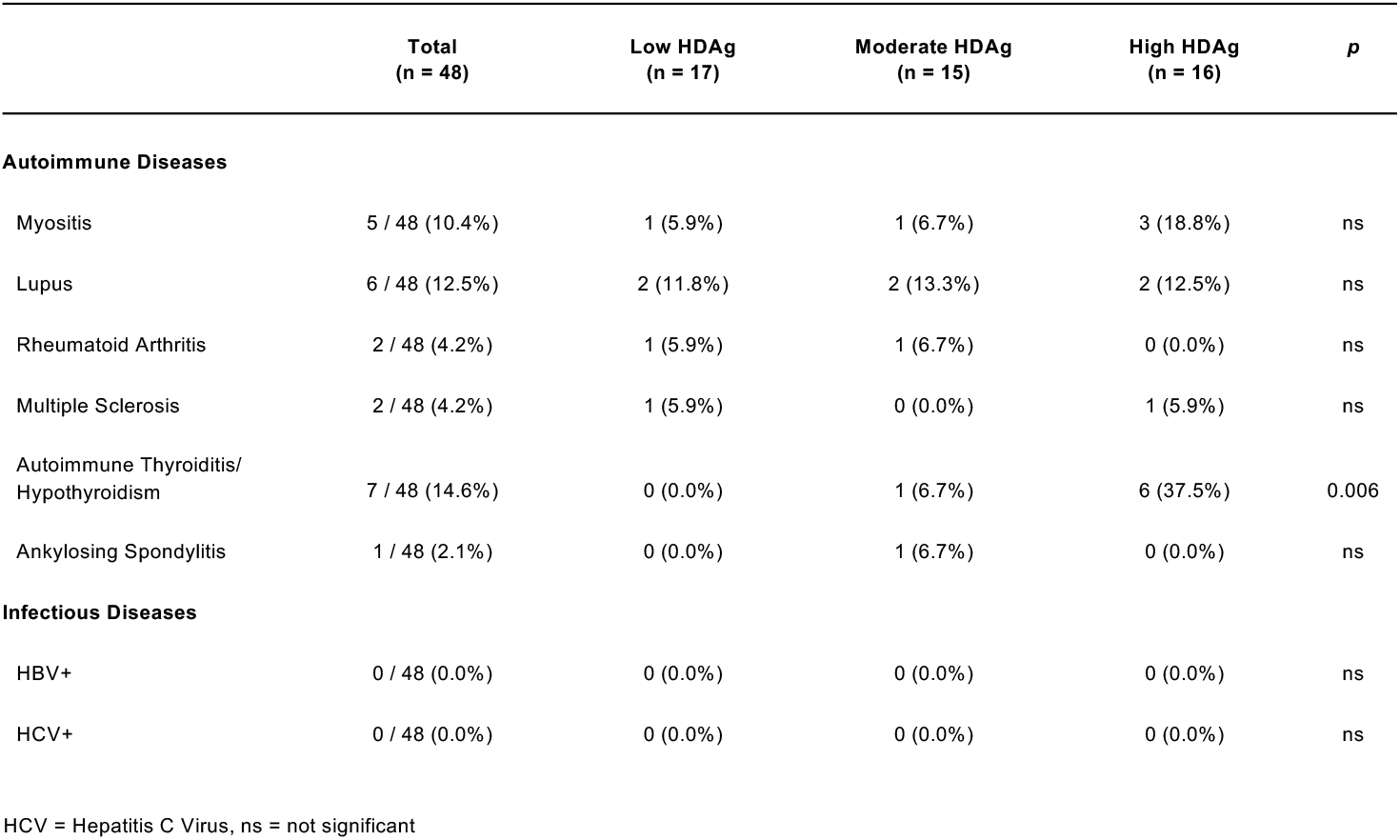
Clinical comorbidities stratified by HDV antigen intensity level.

|  | Total<br>(n = 48) | Low HDAg<br>(n = 17) | Moderate HDAg<br>(n = 15) | High HDAg<br>(n = 16) | p |
| --- | --- | --- | --- | --- | --- |
| **Autoimmune Diseases** |  |  |  |  |  |
| Myositis | 5 / 48 (10.4%) | 1 (5.9%) | 1 (6.7%) | 3 (18.8%) | ns |
| Lupus | 6 / 48 (12.5%) | 2 (11.8%) | 2 (13.3%) | 2 (12.5%) | ns |
| Rheumatoid Arthritis | 2 / 48 (4.2%) | 1 (5.9%) | 1 (6.7%) | 0 (0.0%) | ns |
| Multiple Sclerosis | 2 / 48 (4.2%) | 1 (5.9%) | 0 (0.0%) | 1 (5.9%) | ns |
| Autoimmune Thyroiditis/<br>Hypothyroidism | 7 / 48 (14.6%) | 0 (0.0%) | 1 (6.7%) | 6 (37.5%) | 0.006 |
| Ankylosing Spondylitis | 1 / 48 (2.1%) | 0 (0.0%) | 1 (6.7%) | 0 (0.0%) | ns |
| **Infectious Diseases** |  |  |  |  |  |
| HBV+ | 0 / 48 (0.0%) | 0 (0.0%) | 0 (0.0%) | 0 (0.0%) | ns |
| HCV+ | 0 / 48 (0.0%) | 0 (0.0%) | 0 (0.0%) | 0 (0.0%) | ns |

HCV = Hepatitis C Virus, ns = not significant

### HDV + SjS patients lack markers of liver disease or HBV co-infection

Hepatitis delta virus has been extensively studied in association with an HBV co-infection. Therefore, SjS patient liver enzymes and prior HBV test results were analyzed for a potential current or prior HBV co-infection and undiagnosed liver disease. As detailed in Table 4, 12%, 31%, and 29% of the SjS patients had abnormal AST, ALT, ALP, respectively. There were no significant correlations between abnormal AST, ALT, ALP, or AST/ALT ratio and HDAg intensity (Supplemental Figure 5). Similarly, there were no significant correlations between HDAg intensity and abnormal bilirubin, albumin, or protein levels. Twenty (20) patients had previously undergone testing for HBsAg or HBcAb, and of those tested, no patient had a positive result (Table 4). These markers indicate that the patients in this study show no evidence of an HBV co-infection.

**Table 4.** Clinical characteristics of liver disease and hepatitis markers stratified by HDV antigen intensity level.

|  | Total<br>(n = 42) | Low HDAg<br>(n = 15) | Moderate HDAg<br>(n = 13) | High HDAg<br>(n = 14) | *p* |
| --- | --- | --- | --- | --- | --- |
| **AST U/L** (M, 95% CI) | 38.8 (3.8 - 49.3) | 35.3 (23.0 - 47.6) | 42.5 (15.3 - 69.6) | 39.0 (9.3 - 54.6) | ns |
| Abnormal (%) | 5 (12%) | 2 (13%) | 2 (15%) | 1 (7%) |  |
| **ALT U/L** | 40.2 (29.3 - 51.0) | 37.7 (22.2 - 53.3) | 34.6 (17.1 - 52.1) | 48.0 (3.5 - 71.3) | ns |
| Abnormal | 13 (31%) | 4 (27%) | 3 (23%) | 6 (43%) |  |
| **ALP U/L** | 112.2 (50.6 - 130.9) | 105.9 (95.2 - 116.6) | 102.7 (70.4 - 135.0) | 127.9 (39.5 - 174.3) | ns |
| Abnormal | 12 (29%) | 3 (20%) | 3 (23%) | 6 (43%) |  |
| **Bilirubin mg/dL** | 0.7 (0.3 - 0.8) | 0.8 (0.5 - 1.0) | 0.7 (0.5 - 0.8) | 0.7 (0.4 - 0.8) | ns |
| Abnormal | 4 (10%) | 2 (13%) | 1 (8%) | 1 (7%) |  |
| **Albumin g/dL** | 3.9 (3.5 - 4.1) | 4.0 (3.8 - 4.2) | 3.9 (3.6 - 4.2) | 3.9 (3.5 - 4.1) | ns |
| Abnormal | 6 (14%) | 2 (13%) | 2 (15%) | 2 (14%) |  |
| **Protein g/dL** | 6.8 (6.2 - 7.0) | 6.8 (6.6 - 7.0) | 6.9 (6.4 - 7.4) | 6.8 (6.3 - 7.0) | ns |
| Abnormal | 6 (14%) | 2 (13%) | 2 (15%) | 2 (14%) |  |
| **Tested for HBV** | 20 (48%) | 7 (47%) | 7 (54%) | 6 (43%) |  |
| **HBSAg+** | 0/19 | 0/6 | 0/7 | 0/6 |  |
| **HBcAb+** | 0/15 | 0/5 | 0/5 | 0/5 |  |
| **HBV Diagnosis (ICD-9/10)** | 0 | 0 | 0 | 0 |  |

Abbreviations: M = mean, CI = confidence interval, ALT = alanine aminotransferase, ALP = alkaline phosphatase, AST = aspartate aminotransferase, HBV = hepatitis B virus, HBSAg = hepatitis B surface antigen, HBcAb = hepatitis B core antibody, ns = not significant
\*All patients tested for HBV had a negative test result.

## Discussion

This study details the presence of HDV antigen and genomic sequence within SjS-affected salivary gland tissue. Localization of HDV antigen was most prominent in acinar and ductal cells within the MSG, but was also observed in connective tissues, including adipocytes. On a subcellular level, HDAg localized within the nuclei, cytoplasm and mitochondria. This study demonstrated the first localization of HDV RNA sequence within human MSG tissue and further substantiates the presence of HDV signatures in non-hepatic tissues. The identification of HDV antigen and sequence in SjS-affected MSG with documented capacity to trigger a SjS symptomology *in vivo* (11) further supports a viral-mediated mechanism contributing to SjS development.

Comparative analysis identified multiple clinical features correlating with HDAg expression. The intensity of HDAg expressed in MSG negatively correlated with focal inflammation. A steady decreasing trend of HDAg intensity across SjS patient age groups was also observed (Figure 4C-D). Together, these data demonstrate HDAg expression is elevated in younger patients and in patients with less focal inflammation within MSG tissue. These observations may suggest inflammation within MSG may hinder HDAg expression, favor clearance of HDAg, or SjS-associated inflammation may be independent of HDAg expression. The latter is supported by the lack of detectible antibodies to HDV antigen in HDV+SjS patients (11). Additionally, the higher level of HDAg in younger SjS patients could be due to changes in HDAg expression during SjS development and progression. Prior studies have reported increased MSG inflammation in older SjS patients (27). The question remains as to whether the SjS-associated inflammation is favoring HDAg clearance or if other factors are playing a role in decreased HDAg expression with increased inflammation and age.

Co-localization of HDAg with mitochondria has not been reported prior to this study. Casaca et al demonstrated that HDAg interacts with multiple mitochondrial-sourced genes, including ND2, ND4, COX I, COX II, ATP8, and a mitochondrial-localized protein (NDUFB7) in a yeast two-hybrid screening (28). This would suggest that HDAg may localize to the mitochondria to interact with mitochondrial proteins. Anti-mitochondrial antibodies (AMA) occur in 1.7-13% of Sjogren’s syndrome patients and most often in patients with primary biliary cirrhosis (PBC) (29,30). Increased prevalence of anti-mitochondrial antibodies (AMA) has also been reported in PBC (31), but no direct associations between detectible AMA and HDV have been reported. The detected co-localization of HDAg with mitochondria and its potential impact on mitochondrial function and cellular respiration warrants further investigation.

Autoimmune thyroiditis and hypothyroidism were increased in patients with elevated HDV antigen expression (Table 3). This is in line with prior studies reporting an increase in autoimmune thyroid disease (AITD) diagnoses in patients with SjS (32,33). Caramaschi et. al. noted SjS patients with thyroid disease had a less pronounced SjS phenotype (34). This is consistent with our findings of decreased foci score and autoantibody profiles in patients with elevated HDV antigen expression. Lu et. al. noted an increased risk of hypothyroidism diagnosis in younger (20-44 years of age) SjS patients (35). A similar trend was observed in our analysis with increased HDAg intensity in younger age groups. The question remains as to whether the SjS and AITD share a common pathogenesis and whether HDV is involved. Based on these findings, large scale studies are warranted to further characterize the link between HDV in SjS and risk of other autoimmune diseases and potential shared underlying etiologies.

Multiple limitations must be taken into consideration in interpreting the results of this study. First, this was a retrospective study which relied on testing and diagnoses performed through the standard of care and not part of a prospective clinical study. This approach may have been impacted by the incomplete testing and diagnoses performed across the cohort analyzed. Second, the small study size may impact the strength of observed in situ and clinical feature associations with the HDV profile. Minor salivary gland biopsies are not regularly performed in the diagnosis of SjS and thereby limit the cohort size. Large scale clinical studies are required to further define the in situ and clinical feature associations between salivary gland-localized HDV and the SjS phenotype. To support large scale evaluation of HDV in the SjS patient population, increased utilization of the MSG biopsy for SjS diagnosis would be needed and/or the identification of systemic markers for the HDV(+)/HBV(-) profile would need to be identified and validated.

The question remains as to how SjS patients are acquiring HDV within salivary gland tissue in the absence of a detectible current or past HBV co-infection. Classically, HDV is defined as requiring HBV for packaging and transmission but can actively replicate and persist in the absence of an active HBV co-infection through utilization of host cellular machinery (36). In this retrospective study and in our prior cohort study (11), patients with detectible HDAg expression in MSG tissue lacked evidence of a current or past HBV co-infection or strong markers of viral hepatitis (Table 4, Supplemental Figure 5). Multiple studies have evaluated HBV as a potential trigger for SjS and have concluded a similar or lower incidence of HBV in SjS compared to the general population (37–40). Recent reports have detected HDV in a limited number of HBV-/HCV+ patients (41). None of the SjS patients evaluated in this study had evidence of a current or past HCV infection (Table 3). Advanced studies are required to evaluate alternative mechanisms of HDV localization to salivary gland tissue and in the absence of evident current or past HBV co-infection.

Our knowledge of the hepatitis delta virus is rapidly evolving. Recently, HDV was classified under the Kolmioviridae family (42). This was spurred by the discovery of at least 8 new HDV-like sequences isolated from reservoirs across the animal kingdom and in the absence of readily detectible HBV or other Hepadnaviridae co-infections (43–48). Additionally, HDV has demonstrated the capacity to package into non-HBV particles *in vitro* including vesiculovirus, flavivirus, hepacivirus, arenaviruses and orthohantaviruses (49,50). Currently, there are limited reports of detected HDV without evident HBV coinfections in vivo and in humans (11,41,51). These recent discoveries suggest that HDV and HDV-like sequences may have the capacity to package into non-HBV particles and shift HDV tropism to non-hepatic tissues.

New therapies are being evaluated for the treatment of HDV (21). Bulevirtide is a new treatment of HBV and HBV/HDV co-infections and acts by blocking the uptake of HBV mediated by the NCTP receptor (52). Lonafarnib is a farnesyl transferase inhibitor and blocks the post translational modification of the large HDAg that is utilized to package the HDV ribonucleoprotein complex into HBSAg coated particles (53,54). While both of these drugs are showing promise in the treatment of HBV/HDV co-infections (21,55,56), neither directly target HDV replication or persistence and both rely on an active HBV co-infection. Prior studies have evaluated the potential for HDAg-targeted vaccines and demonstrated limited efficacy in targeting HDV in HBV or woodchuck hepatitis virus (WHV) co-infection models (57). It is unclear whether HDAg-targeted vaccination would work differently in an SjS model compared to hepatic localized HDV co-infections. Our prior studies have demonstrated the lack of detectible HDV-Ab in SjS patients with demonstrated HDV antigen and sequence within salivary gland tissue (11). Advanced studies are required to identify viable approaches to target HDV clearance in SjS-affected salivary gland tissue and other potential systemic reservoirs.

Advanced studies are essential to further clarify the role of HDV in Sjogren’s syndrome. It remains unclear how, mechanistically, HDV is triggering the SjS-phenotype within salivary gland tissue. Additionally, larger studies are required to define how HDV is localizing and persisting within SjS salivary gland tissue and how SjS patients are acquiring the unique HDV+/HBV-profile.

## Methods

### Study Design

The study was approved and completed in accordance with the Institutional Review Board (IRB) program coordinated by the University of Utah’s Office of Research Integrity and Compliance (ORIC) (IRB 00098495, “The Role of Hepatitis Delta Virus in Sjogren’s Syndrome”). Diagnostic information was obtained for patients with SjS International Classification of Diseases (ICD9, ICD-10, and associated subcodes) diagnostic code (ICD-9:710.2, ICD-10:M35.0-M35.09) and that had undergone a MSG biopsy. Of these patients, 48 available formalin fixed paraffin embedded (FFPE) labial MSG biopsy blocks acquired between 2014-2021 were identified and utilized in this study. Associated electronic medical records data were analyzed, including patient demographics (age at SjS diagnosis, gender, race/ethnicity), Sjogren’s syndrome (ICD-9: 710.2, ICD-10:M35.0-M35.09) diagnostic codes, associated autoimmune (Ankylosing Spondylitis (ICD-9:720, ICD-10:M45), Autoimmune Thyroiditis (ICD-9:245.2:,ICD-10:E06.3), Hypothyroidism (ICD-9:244.9, ICD-10:E03.9), Multiple Sclerosis (ICD-9:340, ICD-10:G35), Myositis (ICD-9: 729.1,ICD-10:M60), Rheumatoid Arthritis (ICD-9:714, ICD-10:M05), Systemic Lupus Erythromatosus, (ICD-9:710.0, ICD-10: M32)) and infectious disease (Hepatitis B Virus (HBV) (ICD-9:70.2, 70.20, 70.22, 70.3, 70.30, 70.32, ICD-10: B16, B18.0, B18.1), Hepatitis D Virus (HDV) (ICD-9: 70.21, 70.31, 70.42, 70.23, 70.52, 70.33, ICD-10: B16.0, B16.1, B17.0, B18.0) Hepatitis C Virus (HCV) (ICD-9: 70.41, 70.44, 70.51, 70.54, 70.7, ICD-10: B18.2, B19.2) co-morbidities, lab test results (anti-nuclear antibodies (ANA), anti-SSA/Ro-52 and -60, anti-SSB/la, rheumatoid factor (RF), aspartate aminotransferase (AST), alanine aminotransferase (ALT), albumin, bilirubin, alkaline phosphatase (ALP), protein (total), HBsAg (hepatitis B surface antigen), and HBcAb (hepatitis B core antibody)). Focus score for each biopsy was calculated according to previously used methodology and reported as focal lymphocytic infilitrates (≥50 lymphocytes) per 4mm^2^ area (58). Percentage of adipose tissue and focus score in tissue sections were measured on hematoxylin and eosin-stained slides.

### Immunohistochemical Analyses

Immunohistochemical staining of FFPE human salivary gland tissue was conducted using citrate buffer (10mM sodium citrate, pH6.0, 0.05% Tween20) for antigen retrieval, followed by 1 hour incubation in blocking solution (3% BSA in TBS) at room temperature. Incubation with primary antibodies was performed overnight at 4°C. The following primary antibodies were used in staining procedures: rabbit monoclonal anti-HDAg antibody (1:400; gift from John Taylor, PhD), mouse monoclonal anti-Aquaporin 5 (APQ-5) antibody (1:400; Santa Cruz Biotechnology Inc, SC-514022), mouse monoclonal anti-Cytokeratin 7 (CK-7) antibody (1:200; Abcam, ab9021), and OxPhos antibody cocktail (1:200; Thermo Fisher Scientific, #458199). Anti-HDAg was detected using donkey anti-rabbit IgG (H+L) Alexa Fluor® 488 conjugate (1:500; Invitrogen, A21206), and Anti-AQP5, Anti-CK7, and OxPhos antibody cocktail were each detected using goat anti-mouse IgG, (H+L) Alexa Fluor® 594 conjugate (1:500, Invitrogen, A11032). Labeled secondary antibody incubation was performed for 1 hour at room temperature after which the slides were mounted using Fluoromount-G Mounting Medium with DAPI counterstain (Invitrogen, #00495952). A Nikon Eclipse Ti2-S-HW microscope was used for all imaging and NIS-Elements AR 5.11.00 (Nikon, Japan) software was used for quantitative assessment of HDAg in tissue samples.

### HDAg Intensity Analysis

HDAg expression intensity was determined through intensity sampling of 8 separate 0.15mm² sections of each tissue sample using NIS-elements AR 5.11.00 (Nikon, Japan). HDAg intensities were employed as a comparative measure of HDAg expression levels in our analyses. Patient samples were divided into three groups based on HDAg intensity level. “High” and “low” intensity groups included patients with HDAg intensity levels above or below the upper and lower limits of the confidence interval for the HDAg intensity median of the cohort. “Moderate” intensity group was defined as patient samples that were within the 95% confidence interval for the median HDAg intensity of the cohort.

### I*n situ* Hybridization of HDV RNA

Custom probes were designed and ordered from Advanced Cell Diagnostics to detect sense HDV RNA, anti-sense HDV RNA, and mitochondrial COX I RNA (ACD, 546811-C3). *In situ* hybridization was conducted according to the RNAscope® protocol for formalin-fixed, paraffin-embedded tissue samples. Specimens were processed per the manufacturer’s protocols. Briefly, deparaffinized and dehydrated samples were then treated with RNAscope® Hydrogen Peroxide for 10 min. Tissue sections were incubated in boiling 1X Target Retrieval reagent (100°C to 103°C) for 15 min, followed by incubation in 100% ethanol, and then dried. Tissue sections were then treated with RNAscope® Protease Plus at 40°C for 30 min. The slides were then incubated with custom probe mixes at 40°C for 2 hrs and washed with 1X RNAscope® Wash Buffer. Slides were stored overnight at room temperature in 5X saline sodium citrate (SSC). Sections were treated the following day with manufacturer-supplied AMP1 and AMP2 reagents, each for 30 minutes, followed by treatment with AMP 3 reagent for 15 min; washes with 1X wash buffer were conducted between each incubation. Slides were then incubated for 15 min at 40°C with their respective HRP channel reagent dependent upon which probes were applied to the tissue (sense HDV, anti-sense HDV, or COX I). Slides were washed with 1X Wash Buffer and then incubated with their respective fluorophore for 30 min at 40°C. The following fluorophores were used during incubations: Opal 520 Reagent Pack (1:1500; Akoya Biosciences, FP1487001KT) for COX I detection, Opal 570 Reagent Pack (1:1500; Akoya Biosciences, FP1488001KT) for detection of HDV RNA. Following incubation with fluorophores, slides were washed with 1X Wash Buffer and incubated with HRP Blocker reagent for 15 min at 40°C. Slides were washed once more with 1X Wash Buffer and mounted using Fluoromount-G Mounting Medium with DAPI counterstain (Invitrogen, #00495952) and imaged. Validation of RNAscope® probe was done using an HEK-293T cell line expressing S-HDAg (Supplemental Figure 3).

## Statistics

Patients’ minor salivary gland adipose tissue, focus scores, age, and associated lab tests were evaluated using linear regression analysis and ANOVA. Welch’s T-test was used to evaluate HDAg intensity and focus score differences based on sex and auto-antibody test results. A Fisher’s exact test was utilized to assess statistical differences in demographics, age groups, and comorbidities based on patients’ HDAg intensity group. GraphPad Prism software (9.4.1) was used for statistical analyses with a p-value ≤ 0.05 considered significant.

## Author Contributions

MH and MW contributed to research design, literature review, IRB approval, lab-based experiments, data analysis, and manuscript preparation. SF contributed to the literature review, lab experiments, and data analysis. BF contributed to the literature review. All authors provided approval for publication. Author order reflects the relative size and contributions to the project and manuscript.

## Acknowledgments

This study was supported by funding from the National Institute of Health (NIH), the National Institute of Dental and Craniofacial Research (NIDCR R00 DE021745), the University of Utah School of Dentistry, the Undergraduate Research Opportunities Program (UROP) at the University of Utah, and the Summer Program for Undergraduate Research (SPUR) at the University of Utah. Minor salivary gland biopsies and accompanying pathology reports and electronic medical record data requests were obtained from the University of Utah’s UHealth system, UHealth’s dermatopathology and surgical pathology departments and the Data Science Services, respectively. We would additionally like to thank S. Shankar, E. Diggins, B. Acosta, R. Hill for diligent review of this manuscript.

## Supplemental Figures

**Supplemental Table 1**. C3 and C4 Levels by HDAg Intensity Group.

**Supplemental Figure 1. Subcellular colocalization of HDAg with nuclei of adipocytes found in minor salivary gland biopsies. A)** Merged image of adipose tissue in MSG with cell nuclei (blue), HDAg (green) and mitochondrial proteins (red). **B)** Bright field image of adipose tissue morphology. Cell nuclei were stained with

DAPI **(C)** anti-HDAg antibody was used for immunostaining of HDAg **(D)**. OxPhos Human Antibody Cocktail was used for identification of mitochondria **(E)**. Scale bar = 50μm

**Supplemental Figure 2. Examples of HDAg intensity groupings used in data analysis**. *In situ* immunohistochemical staining of formalin-fixed paraffin-embedded salivary gland sections. Cell nuclei were stained with DAPI (blue); Anti-HDAg antibody was used for immunostaining of HDAg (green). scale bar = 100μm.

**Supplemental Figure 3. Validation of RNAScope probe for HDV Genomic. I*n situ* hybridization of custom HDV-genomic RNAScope probe targeting HDV genomic sequence was validated using 293T-HEK cells (negative control) and 293T-SAg cells expression S-HDAg (positive control). A)** 293T cells were utilized to evaluate non-specific binding capacity of the custom HDV-genomic RNAScope probe. 293T control cells show no probe hybridization. **B)** 293T-SAg cells contain an integrated S-HDAg expression cassette regulated by a tetracycline operator (ref: Chang 2006). 293T-SAg cells showed hybridization of the custom HDV-genomic RNAscope probe to HDAg RNA localized within the nucleus. Scale bar = 50μm

**Supplemental Figure 4. HDAg intensity and adipose tissue analysis**. Evaluation of HDAg intensity relative to percent adipose tissue within MSG biopsies identified no significant correlation or difference between **A**) HDAg intensity and percent adipose tissue, **B**) percent area of adipose tissue within the three HDAg intensity groups or **C**) percent adipose tissue and age of SjS patients at biopsy. P values were determined with 1-way ANOVA (**B**) or unpaired t-test (**A** and **C**). R values indicate Pearson correlation coefficient and error bars indicate standard deviation.

**Supplemental Figure 5. Lack of systemic markers of liver disease in SjS with elevated HDAg intensity in minor salivary gland tissue**. No significant difference was observed between **A-B**) Aspartate transaminase (AST), **C-D**) alanine aminotransferase (ALT), **E-F**) alkaline phosphatase (ALP) and HDAg intensity levels. Similarly, no significant difference was observed between **G)** AST/ALT, **H**) bilirubin and **I**) albumin levels and HDAg intensity groups. P values were determined with 1-way ANOVA (**B, D, F-I**) or unpaired t-test (**A, C, E**). R values indicate Pearson correlation coefficient and error bars indicate standard deviation.

