## Supplemental Figures for "Clinical and *In Situ* Characterization of Hepatitis Delta Virus in Sjogren’s Syndrome"

**Supplemental Table 1. C3 and C4 Levels by HDAg Intensity Group.**

| | |
|---|---|
| **C3 (n = 21)** | |
| Low | 0 (0.0%) |
| Normal | 19 (90.5%) |
| High | 2 (9.5%) |
| **C4 (n = 22)** | |
| Low | 0 (0.0%) |
| Normal | 18 (81.8%) |
| High | 4 (18.18%) |

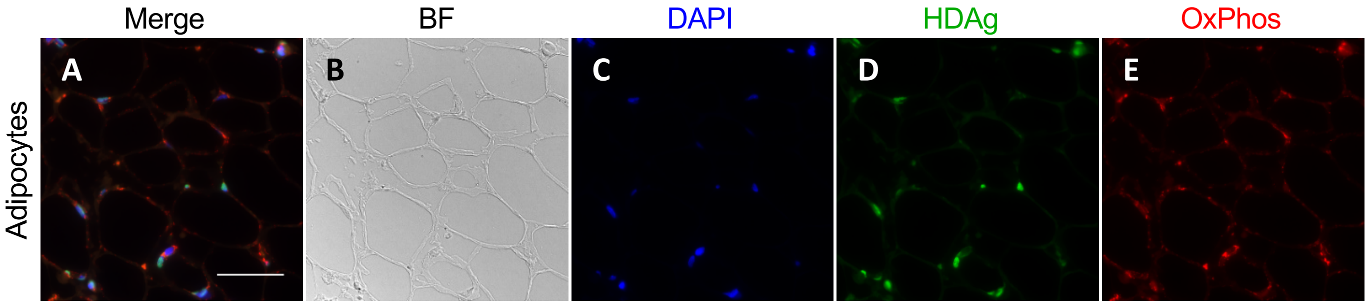


**Supplemental Figure 1. Subcellular colocalization of HDAg with nuclei of adipocytes found in minor salivary gland biopsies. A)** Merged image of adipose tissue in MSG with cell nuclei (blue), HDAg (green) and mitochondrial proteins (red). **B)** Bright field image of adipose tissue morphology. Cell nuclei were stained with DAPI **(C)** anti-HDAg antibody was used for immunostaining of HDAg **(D)**. OxPhos Human Antibody Cocktail was used for identification of mitochondria **(E)**. Scale bar = 50μm

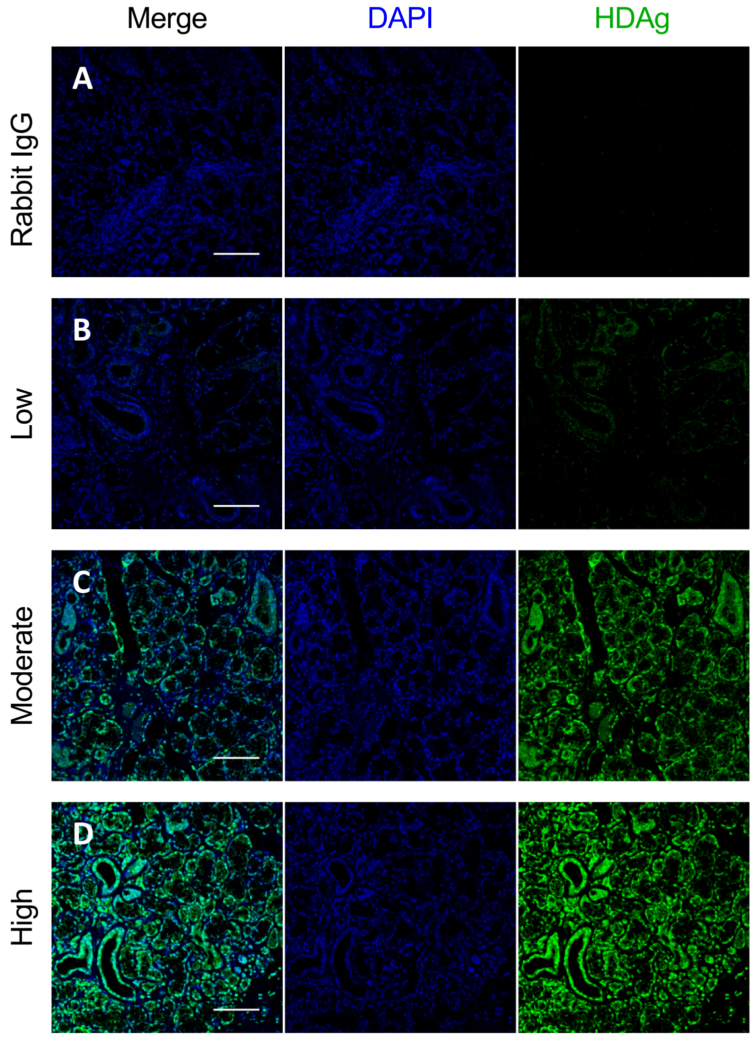


**Supplemental Figure 2. Examples of HDAg intensity groupings used in data analysis.** *In situ* immunohistochemical staining of formalin-fixed paraffin-embedded salivary gland sections. Cell nuclei were stained with DAPI (blue); Anti-HDAg antibody was used for immunostaining of HDAg (green). scale bar = 100μm.

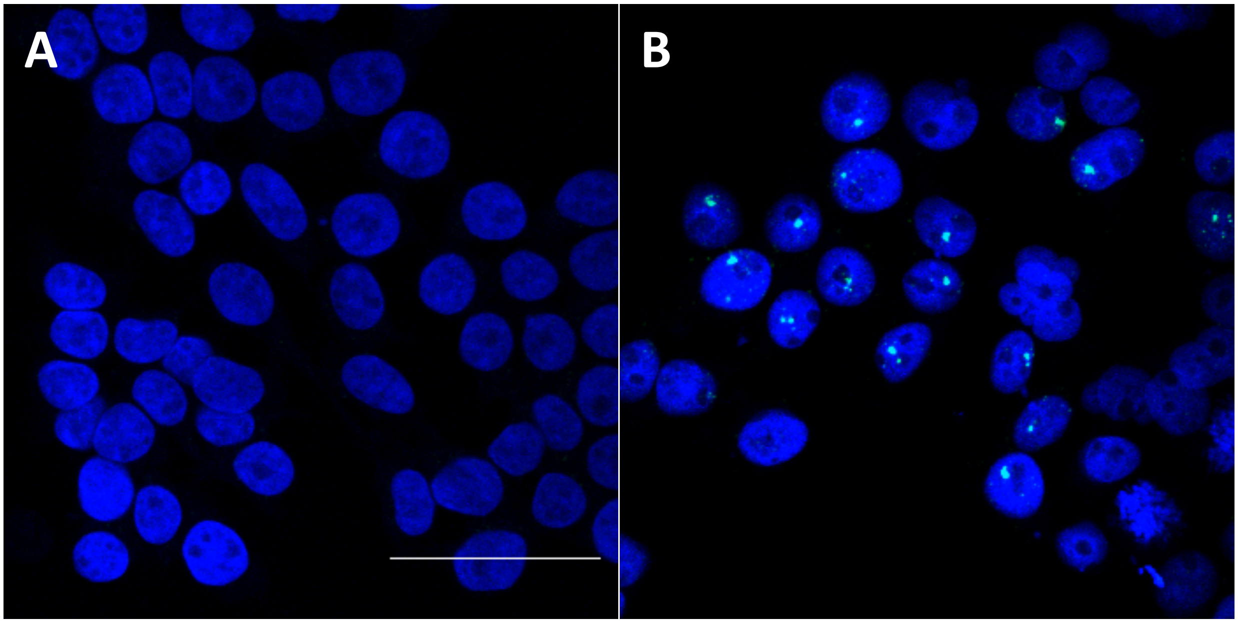


**Supplemental Figure 3. Validation of RNAScope probe for HDV Genomic. I*n situ* hybridization of custom HDV-genomic RNAScope probe targeting HDV genomic sequence was validated using 293T-HEK cells (negative control) and 293T-SAg cells expression S-HDAg (positive control). A)** 293T cells were utilized to evaluate non-specific binding capacity of the custom HDV-genomic RNAScope probe. 293T control cells show no probe hybridization. **B)** 293T-SAg cells contain an integrated S-HDAg expression cassette regulated by a tetracycline operator (36). 293T-SAg cells showed hybridization of the custom HDV-genomic RNAscope probe to HDAg RNA localized within the nucleus. Scale bar = 50µm

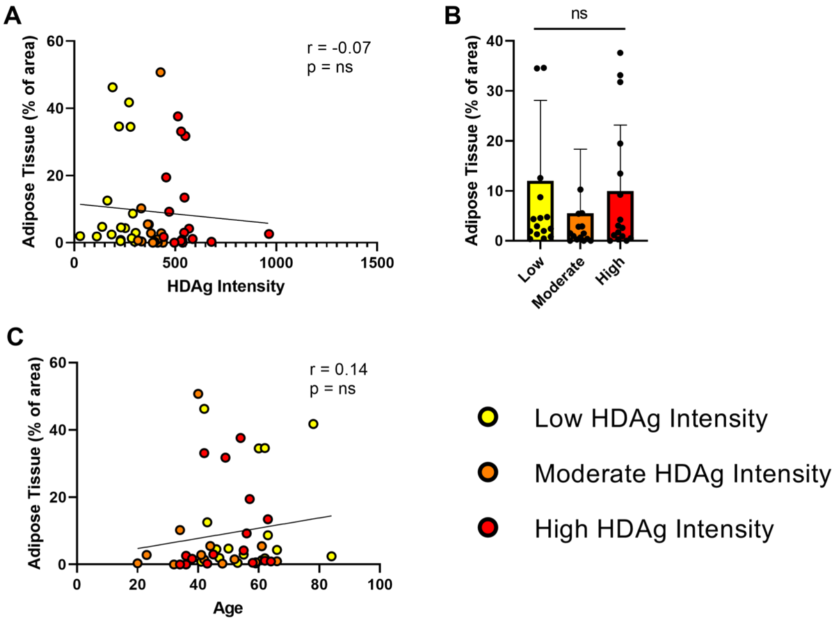


**Supplemental Figure 4. HDAg intensity and adipose tissue analysis.** Evaluation of HDAg intensity relative to percent adipose tissue within MSG biopsies identified no significant correlation or difference between **A**) HDAg intensity and percent adipose tissue, **B**) percent area of adipose tissue within the three HDAg intensity groups or **C**) percent adipose tissue and age of SjS patients at biopsy. P values were determined with 1-way ANOVA (**B**) or unpaired t-test (**A** and **C**). R values indicate Pearson correlation coefficient and error bars indicate standard deviation.

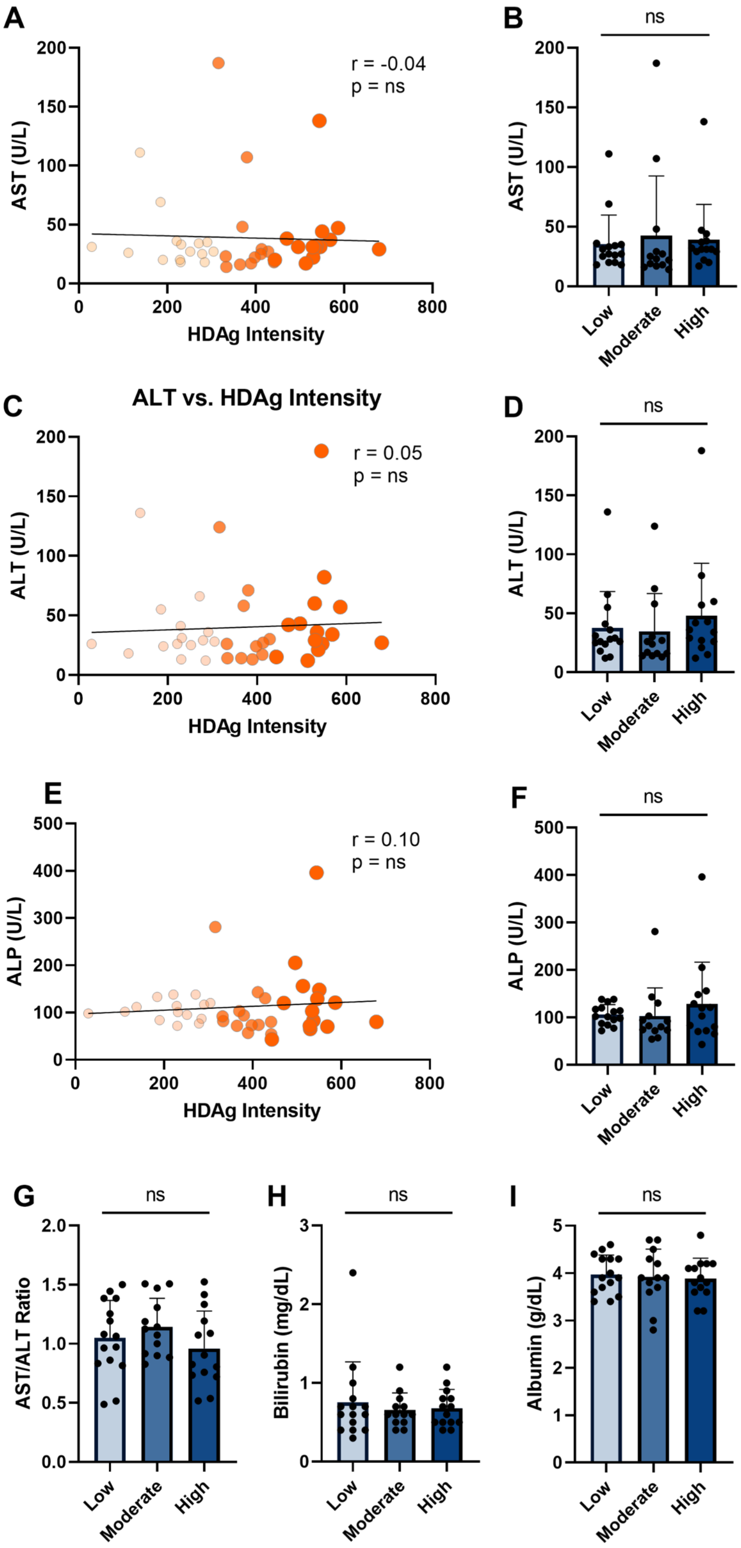


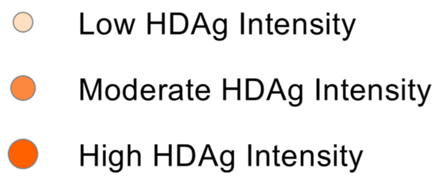


**Supplemental Figure 5. Lack of systemic markers of liver disease in SjS with elevated HDAg intensity in minor salivary gland tissue.** No significant difference was observed between **A-B**) Aspartate transaminase (AST), **C-D**) alanine aminotransferase (ALT), **E-F**) alkaline phosphatase (ALP) and HDAg intensity levels. Similarly, no significant difference was observed between **G)** AST/ALT, **H**) bilirubin and **I**) albumin levels and HDAg intensity groups. P values were determined with 1-way ANOVA (**B, D, F-I**) or unpaired t-test (**A, C, E**). R values indicate Pearson correlation coefficient and error bars indicate standard deviation.
